# A Transcriptomic Hourglass In Brown Algae

**DOI:** 10.1101/2024.04.20.590401

**Authors:** Jaruwatana S. Lotharukpong, Min Zheng, Remy Luthringer, Hajk-Georg Drost, Susana M. Coelho

## Abstract

Complex multicellularity has emerged independently across a few eukaryotic lineages and is often associated with the rise of elaborate, tightly coordinated developmental processes. How multicellularity and development are interconnected in evolution is a major question in biology. The hourglass model of embryonic evolution depicts how developmental processes are conserved during evolution, predicting morphological and molecular divergence in early and late embryo stages, bridged by a conserved mid-embryonic (phylotypic) period linked to the formation of the basic body plan. Initially found in animal embryos, molecular hourglass patterns have recently been proposed for land plants and fungi. However, whether the hourglass pattern is an intrinsic feature of all developmentally complex eukaryotic lineages remains elusive. Here, we tested the prevalence of a (molecular) hourglass in the brown algae, the third most developmentally complex lineage on earth that has evolved multicellularity independently from animals, fungi, and plants. By exploring the evolutionary transcriptome of brown algae with distinct morphological complexities, we uncovered an hourglass pattern during embryogenesis in developmentally complex species. Filamentous algae without a canonical embryogenesis display an evolutionary transcriptome that is most conserved in multicellular stages of the life cycle, whereas unicellular stages are more rapidly evolving. Our findings suggest that transcriptome conservation in brown algae is associated with cell differentiation stages, but not necessarily linked to embryogenesis. Together with previous work in animals, plants and fungi, we provide further evidence for the generality of a developmental hourglass pattern across complex multicellular eukaryotes.

## Introduction

Multicellularity evolved many times during the evolutionary history of eukaryotes (Grosberg and Strathmann 2007). In most cases, the emergence of multicellularity from unicellular ancestors produced relatively simple life forms, but in a small number of lineages this transition was followed by an elaborate series of events that gave rise to what can be considered ‘complex multicellular’ organisms, associated with the evolution of intricate developmental processes (Knoll 2011; Niklas and Newman 2013). The emergence of complex multicellularity is thus thought to be a rare event, having occurred independently in animals, fungi, plants, red and brown algae (Knoll 2011). With the rise of complex multicellular lineages, a major question is how developmental processes can accommodate evolutionary change. In other words, how can the benefits of evolutionary change be co-opted without disrupting intricate developmental processes?

A recurring pattern of evolutionary conservation and variation across developmental stages during the life cycle of multicellular organisms was already observed by 19^th^ century comparative embryologists, who noticed the striking morphological similarity between embryos (von Baer 1828; Müller 1864; Haeckel 1866; His 1875). This observation was more recently revisited at the molecular level, where the morphological pattern of evolutionary conservation and variation is supported by an analogous pattern at the transcriptomic level (Drost et al. 2017; Yanai 2018; Richardson 2022). Two models have been proposed to explain how conserved developmental processes accommodate evolutionary change: the *early conservation model* and the *developmental hourglass model* (and hybrids between both models). According to the ‘early conservation model’, evolutionary change is increasingly permitted as embryo development proceeds, which presents a low-mid-high pattern of evolutionary novelty and originates from von Baer’s third law of embryo development (von Baer 1828). In contrast, the ‘developmental hourglass model’ proposes that evolutionary change is restricted in the mid-phase of embryo development, presenting a high-low-high pattern of evolutionary novelty. This model is motivated by morphological differences observed in the early phases of embryogenesis (e.g. diversity in embryo cleavage patterns), similarity in mid-embryogenesis (as embryos converge on a basic body plan) and differences in the later phases (as embryos acquire further species-specific features) (Duboule 1994; Raff 1996).

Using transcriptome novelty as a quantitative readout for evolutionary novelty (evolutionary transcriptomics), some early studies have reported early conservation patterns (Roux and Robinson-Rechavi 2008; Comte et al. 2010; Piasecka et al. 2013), while more recent studies have reported hourglass patterns across various multicellular eukaryotic lineages using bulk (Domazet-Lošo and Tautz 2010; Kalinka et al. 2010; Irie and Kuratani 2011; Quint et al. 2012; Cheng et al. 2015; Levin et al. 2016) and single-cell transcriptomics (Ma and Zheng 2023; Mayshar et al. 2023; Ullrich and Glytnasi 2023). It should be noted though that different biological properties such as pleiotropically expressed genes, mutational robustness, inter-embryo expression variability, chromatin accessibility and enhancer conservation may follow different models (Uesaka et al. 2022). The empirical findings at the transcriptome level further tie in with theoretical modelling, which supports the narrative of the natural emergence of hourglass-like structures in complex evolving systems (Akhshabi et al. 2014; Friedlander et al. 2015; Sabrin and Dovrolis 2017).

However, our understanding of the generality of the developmental hourglass across complex multicellular eukaryotes is incomplete without considering the brown algae. Brown algae, or brown seaweeds, belong to the stramenopiles, a large supergroup of organisms that are only distantly related to animals, land plants and fungi (Bringloe et al. 2020). Notably, brown algae have independently evolved complex multicellularity by 450 Ma (Choi et al. 2024), and have since become the third most morphologically complex lineage on earth, rivalling plants in terms of sophistication in developmental patterns (Bringloe et al. 2020). In addition, brown algal species present a wide range of morphological and developmental complexity. For example, the ‘simple’ filamentous *Ectocarpus* is composed of up to eight cell types (Coelho et al. 2020), and is capable of multicellular growth and differentiation in absence of canonical embryogenesis (Bothwell et al. 2010). In contrast, the giant kelps (Laminariales) and Fucales undergo obligatory, canonical embryogenesis to generate meters-long adult individuals composed of dozens of cell types (Bringloe et al. 2020). By harnessing the diversity of developmental complexity in brown algae, we can disentangle the effect of multicellular complexity *per se* versus embryogenesis on the evolutionary transcriptome. Furthermore, sexual systems also evolved independently in brown algae (Barrera-Redondo et al. 2024). During the course of sexual development in brown algae, most brown algae (e.g. *Ectocarpus*) alternate between haploid (gametophyte) and diploid (sporophyte) generations, each consisting of morphologically distinct, complex multicellular forms connected by three different unicellular stages: gametes, meiospores and mitospores (Coelho and Cock 2020; Heesch et al. 2021) (**Fig. 1**). We can thus distinguish the potential role of selection in gametes (e.g. due to sperm competition) from unicellularity (i.e., bottlenecks during the life cycle) (Godfrey-Smith 2016). Alongside the convergent evolution of complex multicellularity, these lineage-specific features make brown algae a unique and powerful system to distinguish overlapping processes seen in animal and plant development.

**Figure 1.**
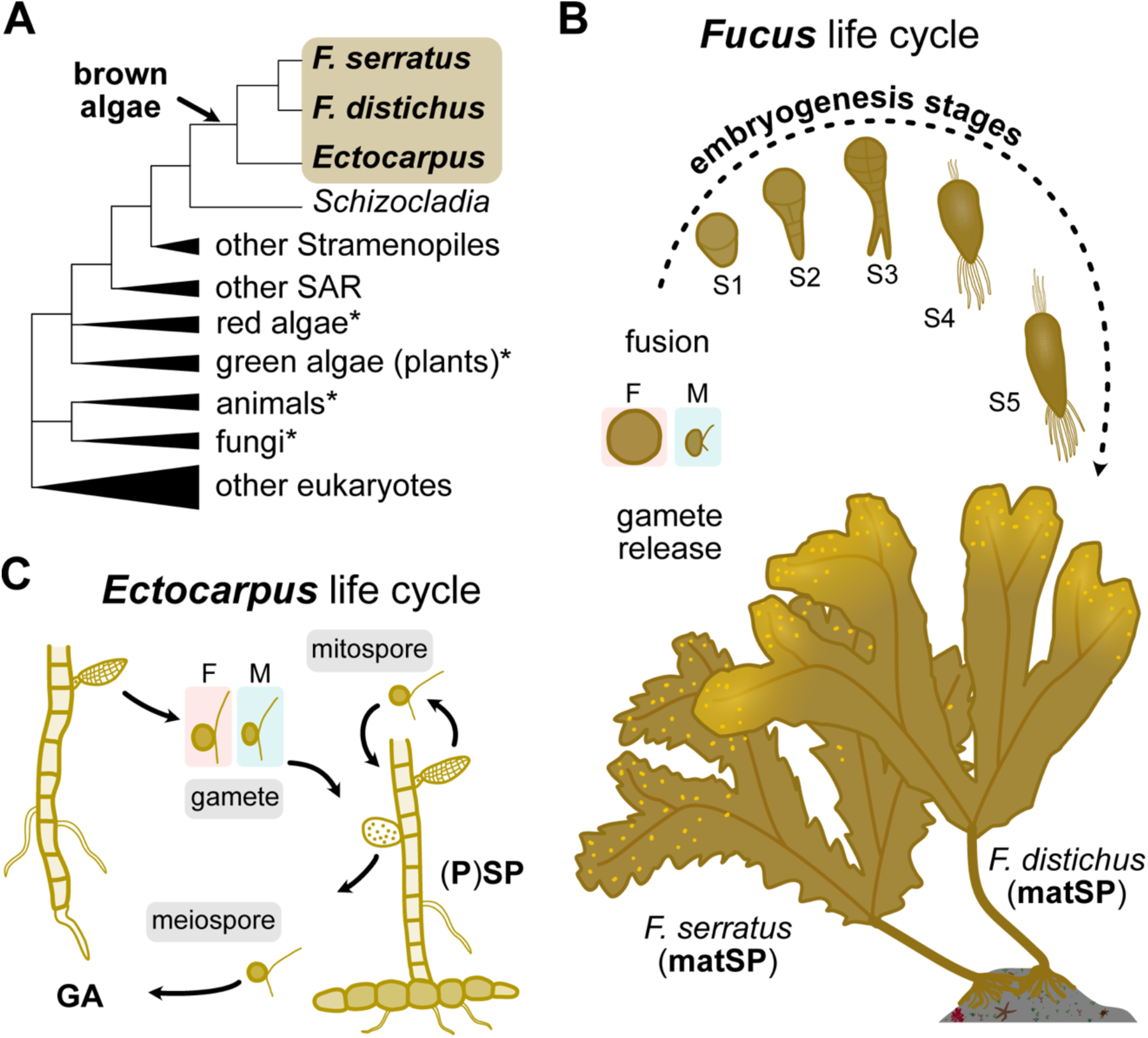
Developmental and morphological diversity in brown algae. (A) Phylogenetic position of *F. serratus*, *F. distichus* and *Ectocarpus* in a simplified eukaryotic tree of life. An arrow marks the independent origin of complex multicellularity in brown algae. The asterisks mark other lineages that evolved complex multicellularity. (B) The life cycle of *Fucus* species. (C) The life cycle of *Ectocarpus*. Unicellular stages are highlighted in grey. GA denotes multicellular gametophytes and (P)SP denotes both multicellular sporophytes and morphologically identical partheno-sporophytes.

Leaning on this unique natural history, we explore brown algal species exhibiting distinct levels of multicellular complexity, to investigate the existence of a developmental hourglass pattern in the third most morphologically complex eukaryotic lineage. If a molecular hourglass pattern does shape the embryo evolution of developmentally complex brown algae, a question arises of whether the same hourglass model underlies the development of all complex multicellular life.

## Results

To test whether a molecular hourglass pattern is underlying brown algal development, we quantified gene expression levels across key ontogenetic stages of development for three distinct brown algal species, *Fucus serratus, Fucus distichus,* and *Ectocarpus sp*. We selected these species to cover the broad diversity of morphological complexity in this group of eukaryotes (**Fig. 1A**). The Fucales are multicellular complex seaweeds with a well-described embryogenesis that occurs highly synchronously without contamination from parental tissues (Goodner and Quatrano 1993). *Fucus serratus* has separate male and female sexes, whereas *Fucus distichus* is a co-sexual species, i.e., the same individual produces male and female reproductive structures (**Fig. 1B**). As a comparative model, we used the filamentous brown alga, *Ectocarpus,* which alternates between two simple but morphological distinct forms, the gametophyte and sporophyte, and presents a range of uni- and multicellular stages but not necessarily a canonical embryogenesis (Coelho et al. 2020) (**Fig. 1C**).

### Hourglass-shaped transcriptome conservation profiles during *Fucus* embryo development

We used an evolutionary transcriptomics approach to test the developmental hourglass hypothesis for the early development of the two *Fucus* species. We first collected stage-specific RNAseq data (**Table S1**) and assigned phylogenetic ages to each protein-coding gene using GenEra (Barrera-Redondo et al. 2023) (**Fig. S1**). Combining both the expression and evolutionary information, we computed the transcriptome age indices (TAI) for each stage, which quantifies the weighted mean of the gene age with its transcript expression level (Domazet-Lošo and Tautz 2010; Drost et al. 2018). In total, we captured the evolutionary and expression data for 8,291 genes in *F. serratus* and 7,907 genes in *F. distichus* (Methods, **Table S2**).

TAI profiles across embryonic development revealed a transcriptomic hourglass pattern in both *Fucus* species (**Fig. 2A, B**). The TAI profiles were statistically significant and robust to all RNA-seq data transformations (**Fig. S2**). We further tested the robustness of the observed hourglass patterns by removing genes with ‘noisy’ expression profiles using noisyR (Moutsopoulos et al. 2021) and confirmed that resulting hourglass patterns in both species remained largely significant (**Fig. S3**). Despite a shift in the timing of the developmental stages between the two species, the transcriptome of the most conserved stage of *F. distichus* was also the one most similar to the *F. serratus* stage with highest degree of conservation (**Fig. S4**). Together, these results provide strong evidence that analogous to animals, plants, and fungi, a developmental hourglass pattern is also shaping embryogenesis of both *Fucus* species.

**Figure 2.**
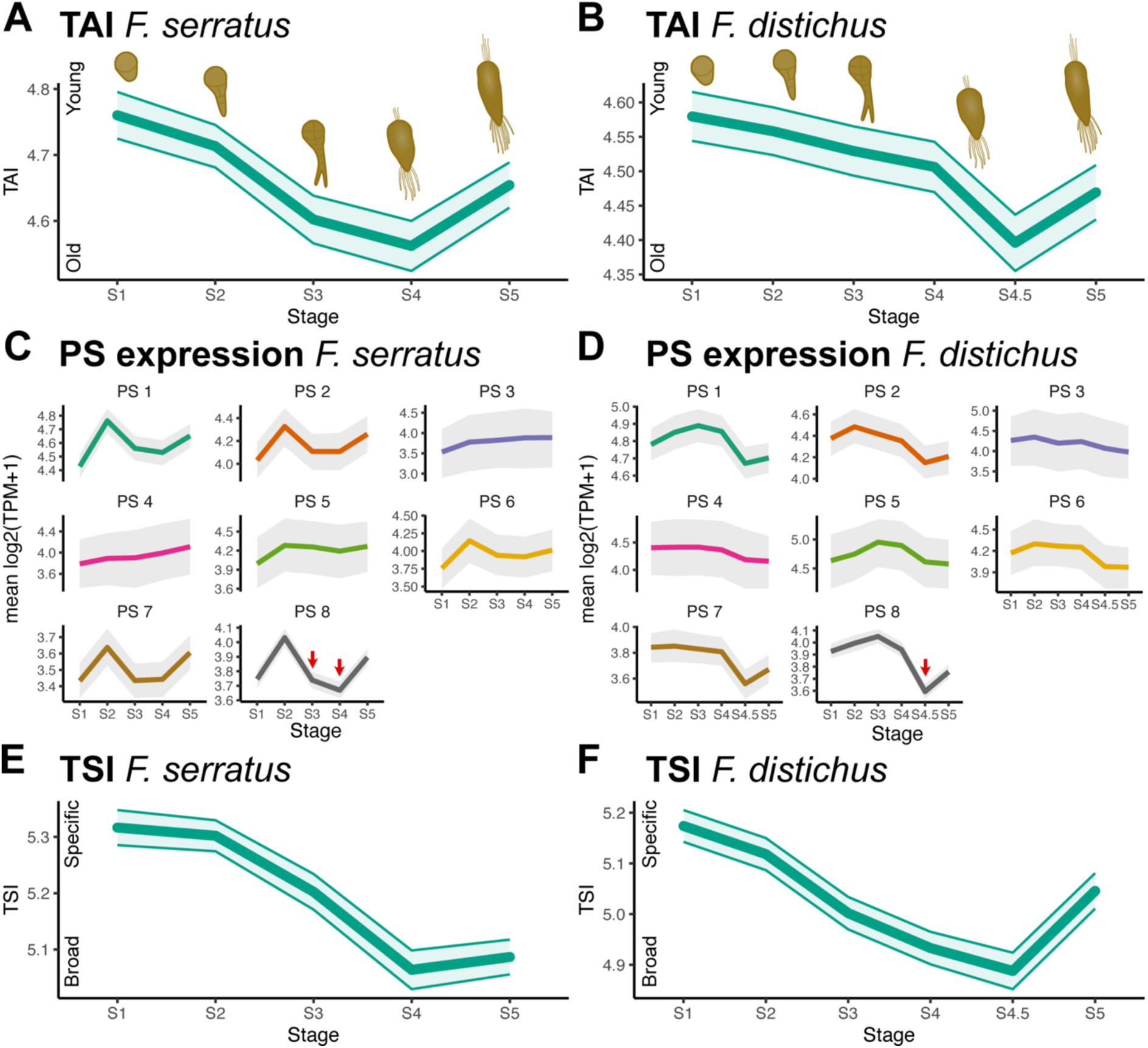
Evolutionary transcriptomes reveal the presence of molecular hourglass patterns in *Fucus* embryogenesis. TAI profile across embryo stages in (A) *F. serratus* and (B) *F. distichus*. A lower TAI marks an older transcriptome (i.e., composed of evolutionarily older genes) and *vice versa*. The mean expression profile of each phylostratum (PS) across embryo stages in (C) *F. serratus* and (D) *F. distichus*. Note, expression is restricted in the evolutionarily youngest genes (PS 8) in both species, as indicated by red arrows. TSI profile of embryogenesis in *F. serratus* (E) and *F. distichus* (F). A lower TSI marks a transcriptome composed of more broadly expressed (temporally pleiotropic) genes and *vice versa*. 50,000 bootstraps were used to compute the standard deviation in (A, B, E, F).

Interestingly, we noticed that evolutionarily young genes (genes in phylostrata [PS] 7 and 8) were markedly downregulated during the waist of the hourglass, rather than older genes being upregulated at this stage (**Fig. 2C, D)**. Therefore, the repression of the young gene expression may be underlying the waist of the hourglass, recapitulating observations in other systems (Quint et al. 2012; Cheng et al. 2015; Drost et al. 2016).

It has been proposed that the conserved transcriptome composition in the waist of the hourglass is caused by higher pleiotropy of genes expressed in these stages (Hu et al. 2017; Liu and Robinson-Rechavi 2018). We examined stage-specificity using *tau* as a proxy for pleiotropy (Yanai et al. 2005; Kryuchkova-Mostacci and Robinson-Rechavi 2017), and computed the resulting transcriptome specificity index (TSI) profile across developmental stages. Stages corresponding to the waist of the hourglass (S4 and S4.5 for *F. serratus* and *F. distichus*, respectively) were represented by more broadly expressed genes (i.e., low *tau*) whereas early and later developmental stages were characterised by more stage-specific genes (**Fig. 2E, F; Table S3**). Therefore, more pleiotropic genes are associated with the evolutionarily conserved stages of development in both *Fucus* species, consistent with findings in animals (Hu et al. 2017; Liu and Robinson-Rechavi 2018).

Finally, we investigated the possible biological processes underlying the evolutionary transcriptomic patterns during *Fucus* early development. Intriguingly, the waist of the hourglass in both *Fucus* species paralleled a major ontogenetic transition, from a ‘cell-type differentiation’ stage, where the algal body plan is established, to a more ‘proliferative’ stage, where development is mainly characterised by somatic cell divisions leading to expansion in the size of the organism (**Fig. S5**). This finding mirrors similar observations in animals and plants, where transcriptomic hourglass patterns mark major ontogenetic transitions (e.g., cell fate acquisition to differentiated cell growth) (Drost et al. 2017).

Together, our observations demonstrate that both *Fucus* species display a transcriptomic hourglass pattern. The conserved mid-embryonic period was characterised by a down-regulation of evolutionarily young genes and an up-regulation of temporally pleiotropic genes, and corresponds with a major developmental transition from cell fate determination to cell proliferation.

### Distinct evolutionary transcriptomes mark *Fucus* adult tissues and sexes

In animals and plants, different tissues and sexes differ in the degree of transcriptome conservation. For example, evolutionarily young genes are disproportionately expressed in the testis of nematodes, flies and mammals (Haerty et al. 2007; Kondo et al. 2017; Villanueva-Cañas et al. 2017; Rödelsperger et al. 2021), and in male reproductive cells in plants (Cui et al. 2015; Gossmann et al. 2016; Julca et al. 2021). To test whether these patterns also characterise sexual differences in brown algae, we examined the TAI across different adult tissues of the two *Fucus* species, as well as between sexes (**Fig. 3A**).

**Figure 3.**
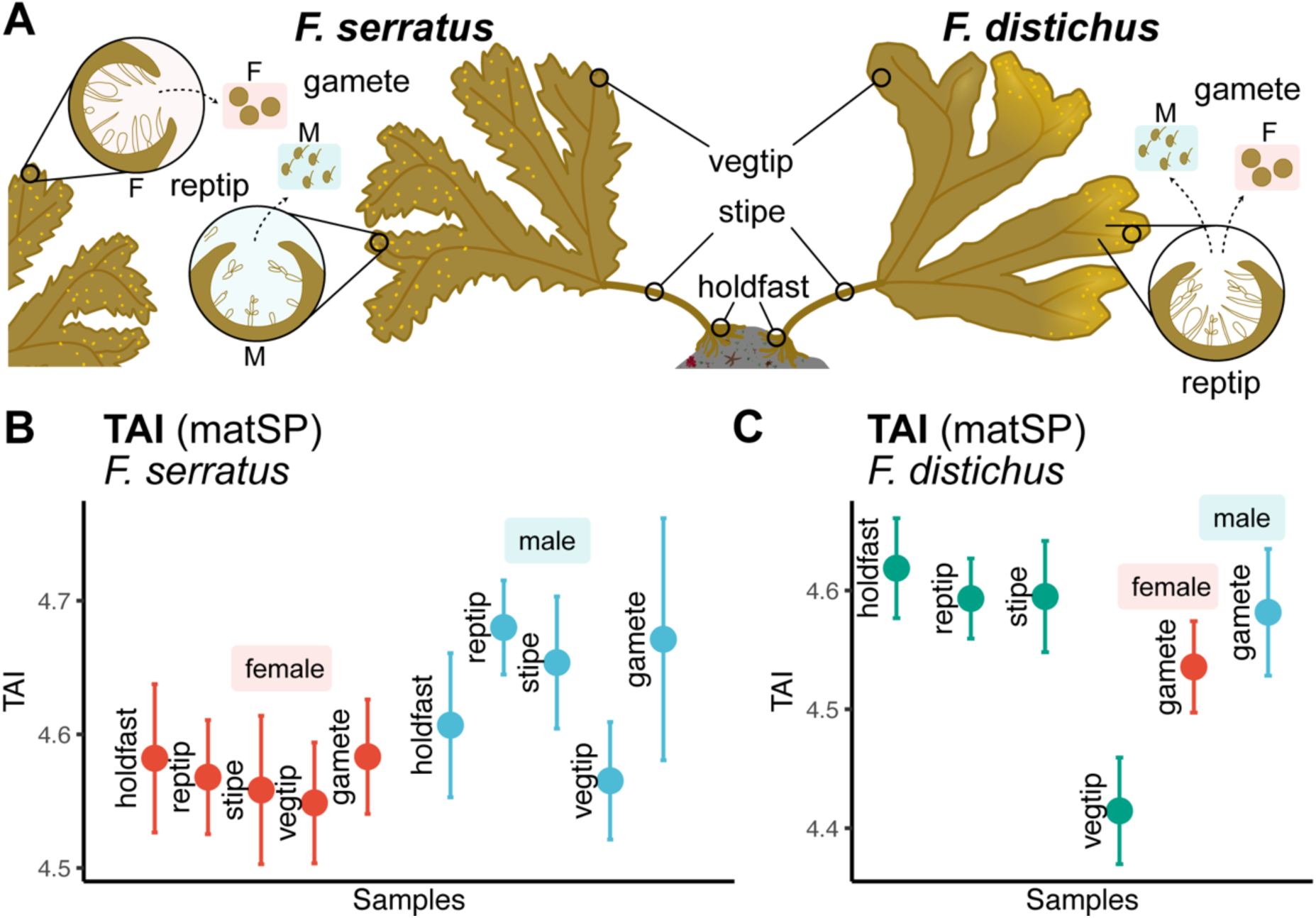
Evolutionary transcriptome landscape across sex and tissues in adult *Fucus*. (A) Outline of adult (mature sporophyte) tissues and gametes in *Fucus* species. Note, *F. serratus* has separate male and female sexes, while the same individual in co-sexual *F. distichus* produces both male and female gametes. (B) TAI profile across adult tissues and gametes in *F. serratus*. (C) TAI profile across adult tissues and gametes in *F. distichus*. A lower TAI marks a transcriptome composed of evolutionarily older genes and *vice versa*. 50,000 bootstraps were used to compute the standard deviation. ‘Vegtip’ and ‘reptip’ denote ‘vegetative tip’ and ‘reproductive tip’, respectively.

In *F. serratus* males, different tissues presented significantly different TAI values (flat line test; p < 0.001), with vegetative tissues consistently exhibiting older transcriptomes compared to reproductive tissues and sperm, which had markedly high TAI **(Fig. 3B**). This difference was also observed in *F. distichus* (**Fig. 3C**). In contrast, TAI values were largely similar across female adult tissues including female gametes (flat line test; p > 0.05) (**Fig. 3B**). We also noticed that sexual differences of TAI were more pronounced at the reproductive tip in *F. serratus* adults, with males having a markedly younger transcriptome compared to females (permutation test; p < 0.001). Intriguingly, the stipe (in both species) also presented relatively young transcriptomes, specifically in males of *F. serratus*. Therefore, similar to animals and plants, *Fucus* males (particularly reproductive tissue and sperm) display an evolutionarily younger transcriptome.

### Multicellularity constrains transcriptome evolution in *Ectocarpus*

In contrast to *Fucus*, *Ectocarpus* morphology is substantially ‘simpler’ (Coelho et al. 2020) but its life cycle comprises two morphologically distinct, free living multicellular generations, the gametophyte and the sporophyte (see **Fig. 1C**), each composed of 3-5 cell types (Cock and Collén 2015). During this alternation of generations, a total of three types of unicell units are produced: gametes, meiospores and mitospores (see **Fig. 1C**), which allows us to disentangle the effect of being a ‘gamete’ *per se* from the effect of being a single cell unit. Furthermore, *Ectocarpus* gametes may develop parthenogenically in absence of fusion, and therefore haploid gametes can develop into adults (partheno-sporophyte) whose morphology closely resembles that of a diploid sporophyte (Bothwell et al. 2010) (see **Fig. 1**). Thus, *Ectocarpus* is a powerful comparative system to test whether the hourglass pattern seen during *Fucus* embryogenesis is the result of constraints imposed by multicellular development *per se* or whether this hourglass pattern of transcriptome conservation is tied to a specific embryogenesis process present in *Fucus* but not in *Ectocarpus*.

We examined the evolutionary transcriptome during the *Ectocarpus* parthenogenetic life cycle, by profiling the transcriptomes of the three unicellular stages (gametes, meiospores and mitospores), three stages during gametophyte development (immature, mature and senescent), and three stages in partheno-sporophyte development (early, immature and mature), for both male and female lines (**Table S1**). As done in *Fucus*, we performed gene age inference (**Fig. S1),** and computed the TAI values from 11,571 genes at each developmental stage (**Table S2**). This analysis revealed that *Ectocarpus* unicellular stages in both males and females exhibited a significantly higher TAI compared to the multicellular stages (**Fig. 4A, S6**), and this pattern was consistent across RNA-seq data transformations (**Fig S7**). In addition, we computed the average purifying selection acting on each stage through the transcriptome divergence index (TDI) metric, where a lower TDI indicates a transcriptome composed of genes under stronger purifying selection and *vice versa* (**Methods, Table S4**). Compared to the TAI data, we noticed that gametes, but not spores, consistently exhibited a high TDI (**Fig. 4B**). Our observations suggest that whilst evolutionarily young genes are more likely expressed in unicellular stages, genes under relaxed purifying selection are disproportionately found in gametes.

**Figure 4.**
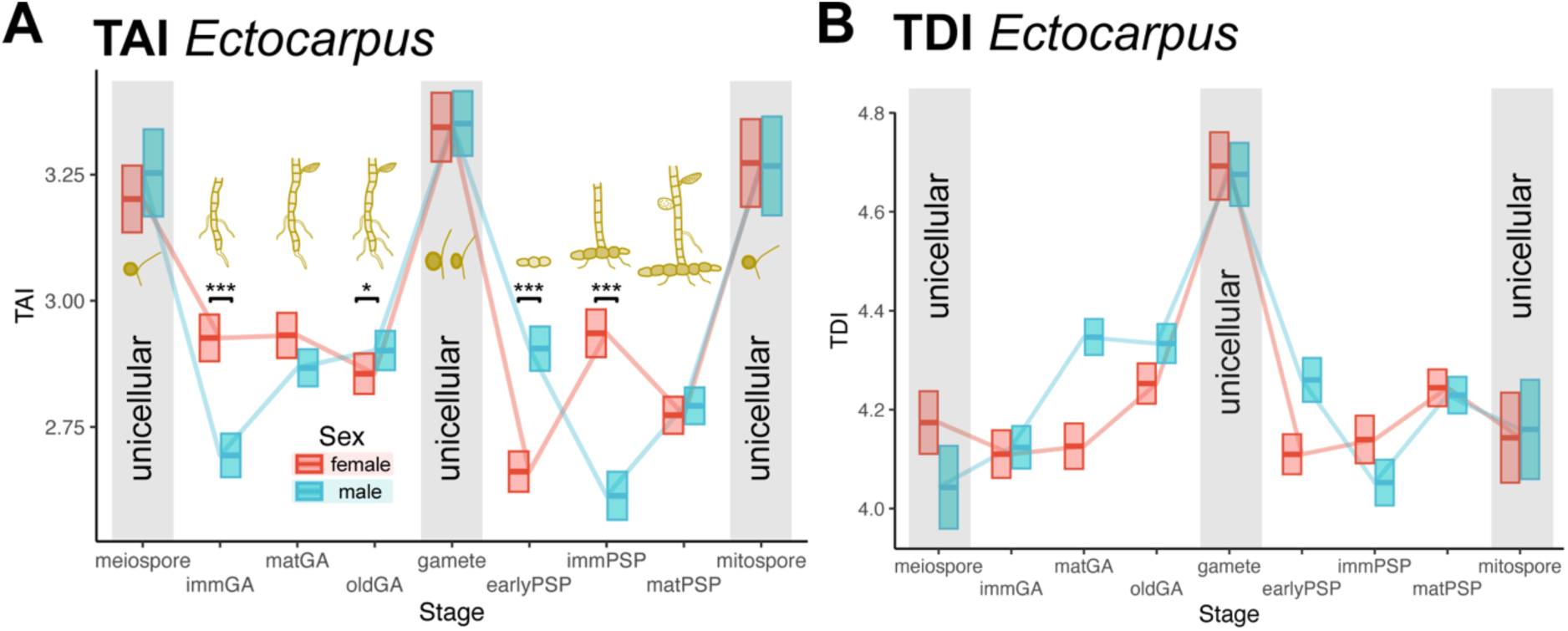
Evolutionary transcriptome dynamics across the life cycle of *Ectocarpus*. (A) TAI profile across *Ectocarpus* life cycle stages. A lower TAI marks a transcriptome composed of evolutionarily older genes and *vice versa*. (B) TDI profile across *Ectocarpus* life cycle stages. A lower TDI marks a transcriptome composed of genes under stronger purifying selection and *vice versa*. Blue lines represent male stages while red lines represent female stages. The upper and lower limits of the boxes demarcate the standard deviation, based on 50,000 bootstraps.

The TAI profile was markedly different between males and females during multicellular development (**Fig. 4A, S6**). *Ectocarpus* males had higher TAI values than females during the early partheno-sporophyte stage, whilst the reverse occurs during the immature gametophyte and partheno-sporophyte stages. Interestingly, the sexual difference in TAI during the gametophyte development culminated at the immature stage, which is the stage when the gametophyte transcriptomes are most different between males and females (Lipinska et al. 2015).

Finally, we focused on the genes that underlie the high TAI values in unicellular stages. Based on the partial TAI value of each individual gene, 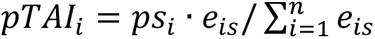 (where e*_is_* denotes the expression level of a given gene *i* at stage *s*, *рs_i_* its gene age assignment, and *n* the total number of genes) (Drost et al. 2018), we identified 115 genes in males and 98 in females that most strongly contributed to the TAI across all unicellular stages (**Fig S8**) (Methods). GO term analysis did not retrieve any functional enrichment, especially since less than 10% of these genes have functional annotation. In contrast, the same analysis using genes with the strongest contribution to the TAI in the multicellular stages returned older genes with GO term enrichment linked to translational, organellar and transcriptional processes in both males and females (**Fig S9**; **Table S5**).

In sum, while in the complex multicellular *Fucus* species, the period of transcriptome conservation was coupled to the transition from cell differentiation to proliferation, in the filamentous *Ectocarpus*, the switch to multicellularity from unicellular, dispersal stages constrains transcriptome evolution.

## Discussion

### Developmental hourglass patterns shape *Fucus* body plan formation

The developmental hourglass model has been reported (and debated) at the molecular level in animals (Roux and Robinson-Rechavi 2008; Comte et al. 2010; Domazet-Lošo and Tautz 2010; Kalinka et al. 2010; Irie and Kuratani 2011; Piasecka et al. 2013; Levin et al. 2016; Uesaka et al. 2022; Ma and Zheng 2023; Mayshar et al. 2023), plants (Quint et al. 2012; Drost et al. 2015; Drost et al. 2016; Gossmann et al. 2016), and fungi (Cheng et al. 2015; Merényi et al. 2022; Xie et al. 2022). Here, we trialled an independently evolved complex multicellular lineage, the brown algae, to examine the prevalence of the transcriptomic hourglass pattern across complex multicellular eukaryotes. We show that embryogenesis of morphologically complex brown algae is underpinned by an evolutionary transcriptomic pattern that is consistent with an hourglass model of embryonic evolution. *Fucus* embryos were more divergent at the earliest and latest stages of embryogenesis but presented a more conserved evolutionary transcriptome during the mid-embryonic period, which serves as a body plan blueprint for the adult organism.

The hourglass pattern in *Fucus* was associated with the pervasive downregulation of evolutionarily young genes, rather than the upregulation of evolutionarily older genes, during the conserved mid-embryonic period, analogous to observations in animals, plants, and fungi (Domazet-Lošo and Tautz 2010; Quint et al. 2012; Cheng et al. 2015; Drost et al. 2016; Ma and Zheng 2023). This pattern also suggests that ancient genes (rather than genes specific to brown algae) form the *Fucus* body plan, pointing to the redeployment of pre-existing molecular toolkit for brown algal complex multicellularity. Moreover, more pleiotropic genes were associated with the evolutionarily conserved stages of development in *Fucus*, mirroring animal embryogenesis (Hu et al. 2017; Liu and Robinson-Rechavi 2018). Importantly, the waist of the hourglass corresponded to a major ontogenetic transition, from a cell-type differentiation stage where the body plan of the adult *Fucus* is established, to a proliferative stage, where development is largely characterised by growth of established cell types. This observation is reminiscent of the transition from *primitive* development to *definitive* development and is consistent with the ‘organisational checkpoint’ hypothesis which postulates that a major transcriptome switch during transitions from cell fate acquisition into multicellular growth phases underlies the transcriptomic hourglass seen in animals, plants, and fungi (Schleip 1929; Sander 1983; Drost et al. 2017).

The evidence we present for the hourglass model in *Fucus*, particularly the evolutionary novelty in the early stages, follows a lineage of studies on the establishment of polarity in brown algal zygotes (Bogaert et al. 2023), which details variability in early embryogenesis. For example, the role of maternal cytoplasmic contribution and extrinsic cues differ markedly between *Fucus* (Kropf et al. 1999; Brownlee et al. 2001), *Dictyota* (Bogaert et al. 2017) and *Saccharina* (Klochkova et al. 2019). These observations are consistent with the variations seen also in early animal and plant development, which is then followed by a more conserved period in mid-embryogenesis (Cridge et al. 2016). We suspect that the hourglass model also describes development in morphologically complex brown algae beyond the transcriptomic level.

### ‘Out of male receptacle’ signatures underlie Fucus sexual development

Evolutionary transcriptomes of adult *Fucus* tissues revealed that male reproductive organs express a younger transcriptome, suggesting that evolutionarily young genes are more permissively expressed. This pattern is likely driven by the presence of male gametes (sperm) in the reproductive organs of male individuals. These observations complement independent findings in animals and plants where young genes are enriched in male reproductive tissue (Vinckenbosch et al. 2006; Gossmann et al. 2016; Murat et al. 2023), indicating convergent evolution of the ‘out of testis’ (or in the case of brown algae, ‘out of male receptacle’) pattern. The younger transcriptome of *Fucus* sperm may be associated with sexual selection, which acts mainly through gamete-level interactions in sessile broadcast spawners such as *Fucus* (Evans and Lymbery 2020). Furthermore, the expression of younger genes in *Fucus* males is consistent with recent findings in brown algae, implicating the female development programme as the more pleiotropic “ground state”, superimposed upon in males (Cossard et al. 2022; Liesner et al. 2024). Thus, in a similar manner to embryogenesis, pleiotropy is linked to transcriptome age.

In contrast, *Ectocarpus* did not exhibit differences in evolutionary transcriptomes in male compared to female gametes. This result is consistent with the low level of sexual dimorphism typical of near-isogamous species (Luthringer et al. 2014; Lipinska et al. 2015), limiting the extent of sexual selection in males compared to females, as well as the smaller set of sex-biased genes in *Ectocarpus* compared to *Fucus* and *Macrocystis* (Hatchett et al. 2023).

### Multicellularity (and not embryogenesis) constrains transcriptome evolution in *Ectocarpus*

Unlike *Fucus*, *Ectocarpus* development consists of two independent, multicellular, morphologically simple, life stages connected by three types of unicellular stages (gametes, meiospores, mitospores). Crucially, the full life cycle can proceed without a canonical embryogenesis (Bothwell et al. 2010). Whilst we did not find a classical ‘hourglass’ signature in *Ectocarpus*, we find that multicellular developmental stages exhibit more conserved transcriptomes compared to unicellular stages. Furthermore, the low TDI associated with multicellular stages of development likely reflects ongoing purifying selection, implicating multicellularity *per se* as under evolutionary constraint. This finding may be due to the lack of a singular ‘mid-embryonic period’ in *Ectocarpus*, where the body plan is established. Instead, *Ectocarpus* exhibits a ‘modular’ development, where organs and tissues are differentiated continuously over time. In other words, the *Ectocarpus* ‘body plan’ needs to be reiterated during multicellular developmental stages. We reason that constraints on morphology are imposed continuously by regulatory complexity that results from dynamic and tissue-specific transcriptional controls, resulting in a conserved transcriptome. Conversely, unicellular stages are more permissive to evolutionary change. Interestingly, while TAI was similarly high for all unicellular stages, only gametes show decreased purifying selection, suggesting a signal for sexual evolution at the sequence substitution level that is specific to gametes. Our results demonstrate that species that lack multicellular organisation via embryogenesis may still exhibit a developmental window with higher transcriptome conservation compared to unicellular stages.

Taken together, we present evidence for the existence of a developmental hourglass pattern in morphologically and developmentally complex brown algae, analogous to hourglass patterns previously reported in animals, plants, and fungi. Given the independent origin of multicellularity in brown algae, this pattern may represent a fundamental characteristic of multicellular complex development across the entire eukaryotic tree of life.

## Methods

### Sample preparation

Details of algal strains used are described in **Table S1**. *Fucus* embryos were prepared as described in (Coelho et al. 2002). Briefly, gametes were allowed to release in 2L natural seawater (NSW), and cleaned using several NSW baths for female gametes and phototaxy for male gametes. Then, gametes were mixed together for an hour and fresh zygotes were cleaned as for female gametes. *F. serratus* and *F. distichus* embryos grow highly synchronously and at least 10,000 developing embryos were flash frozen at specific developmental stages (see **Fig. S5**). Embryos were grown at 14°C in full-strength NSW (*F. serratus*) or at 10°C in half-strength NSW (*F. distichus*) supplemented, for one week, with GeO_2_ (0.4mg/L) to avoid growth of diatoms. Media was changed weekly. For both species of *Fucus*, embryos were grown under neutral days conditions (12h/12h d/night cycle). *Ectocarpus* life cycle stages were grown at 14°C in Provasoli-enriched NSW a 12h/12h d/night cycle and 20 µmol photons m^−2^s^−1^ irradiance, as described previously (Coelho et al. 2012).

Note that *Fucus* and *Ectocarpus* are broadcast spawners, releasing their gametes in the surrounding seawater, where gamete fusion and embryo development takes place. Early development, thus, can be followed in a large number of replicate individual clones that develop highly synchronously, without potentially contaminating parental tissue, greatly facilitating experimental approaches.

### RNA extraction from brown algae

*F. serratus* and *F. distichus* adult tissues were quickly brushed and then rinsed with filtered and autoclaved NSW. Different parts such as holdfast, stipe, meristem, vegetative tissue and reproductive tip were sliced into 0,5 to 1 cm pieces and transferred into 1,5 mL Low-bind Eppendorf tubes. The tubes were snap frozen in liquid N2 and the stored at -80°C until further processing.

The RNA extraction protocol was based on (Cossard et al. 2022; Krasovec et al. 2023). Snap-frozen algae were dry-grinded with a pestle in liquid N2/dry ice and mixed with 750 μL of freshly prepared RNA-extraction buffer (100 mM Tris-HCl pH8,0 (Thermo Fisher AM9856); 1,4 M NaCl; 2% CTAB (Sigma Aldrich, 52365-50G); 20 mM EDTA pH8,0; 1% β*-*mercaptoethanol; 2% Polyvinylpyrrolidon (Thermo Fisher AM9690) preheated to 65°C. Then 250 μL 5M NaCl were added into the tubes. An equal volume of chloroform: isoamylalcohol (24:1) was added and mixed well followed by centrifugation at 10000g for 15 minutes at 4°C. The aqueous phase was removed into RNAse-free tubes and extracted again with 250 μL pure ethanol and an equal volume of chloroform: isoamylalcohol (24:1) as before. RNA was precipitated by adding LiCl (Thermo Fisher, AM9480) to a final concentration of 4M, together with 1% volume of β-Mercaptoethanol, mixing and incubating at -20°C overnight.

The RNA was pelleted by centrifugation at full speed (>18000g) for 45 minutes to 1 hour at 4°C. RNA was washed with 70% cold Ethanol and the pellet was air dried for 3-5 minutes and then the RNA was dissolved in 30 μL RNAse free H_2_O. Residual DNA was eliminated using the TURBO DNase Kit (Thermo Fisher, AM1907) according to the manufacturer’s instructions. The final RNA concentration and size distribution were determined using a Qubit (RNA BR Assay Kit, Invitrogen, Q10210) and an RNA Nano bioanalyzer (Aglient, 5067-1511).

### RNA-seq

The RNA-seq libraries were prepared using commercially available kits according to the manufacturer’s instructions. Poly-A selection for mRNA enrichment was performed using the corresponding NEB kit (#E7490) followed by library preparation using the directional RNA library prep kit from NEB (#E7765). An additional NEB kit Low Input RNA library prep kit (#E6420) was used for samples where large amounts of material were not possible to obtain (meiospores and mitospores) (**Table S1**). The Qubit 1x dsDNA HS assay kit (Invitrogen, Q33230) was used to determine the final DNA concentration of the libraries and the DNA high-sensitive Kit (Aglient, 5067-4626) was employed for bioanalyzer analysis to evaluate the distribution of insert sizes.

Sequencing was performed on a NextSeq2000 instrument with sequencing kit P3-300 (Illumina). The libraries were pooled for sequencing such that for each library we obtained about 30,000,000 reads, corresponding to 9Gb of data (**Table S1**).

RNA-seq datasets were processed using the nf-core/rnaseq pipeline v3.5 (Ewels et al. 2020; Patel et al. 2021). For all three species, expression quantification was performed using salmon v1.5.2 (Patro et al. 2017), to ensure consistency, and imported to R using tximport v1.26.1 (Soneson et al. 2016). For *Ectocarpus*, RNA-seq reads were pseudo-mapped to transcripts inferred for each gene from version 2 of the *Ectocarpus* species 7 genome (Cock et al. 2010). Since public genomes were not available for *F. serratus* and *F. distichus*, the quantification was carried out on recently published *de novo* transcriptome assemblies (Hatchett et al. 2023).

We precluded genes with mean length-scaled TPM (transcripts per million) across samples below 2 from subsequent analyses. In analyses indicated as ‘denoised’, we further removed genes with noise-like behaviour using the ‘counts’ mode of noisyR v1.0.0 (Moutsopoulos et al. 2021).

### Transcriptome age index

The TAI captures the average gene age of a given transcriptome, weighted by the expression level of each gene (Domazet-Lošo and Tautz 2010; Drost et al. 2018). The relative age of each gene in *Ectocarpus*, *F. serratus* and *F. distichus* was inferred using GenEra v1.0 (Barrera-Redondo et al. 2023), based on genomic phylostratigraphy (Domazet-Loso et al. 2007). In brief, GenEra takes all protein coding genes and pairwise aligns these sequences against the taxonomy-resolved NCBI non-redundant database (Sayers et al. 2019; Schoch et al. 2020), using DIAMOND v2.0.14 (‘ultra-sensitive’ mode; e-value < 1e-5) (Buchfink et al. 2021). Next, search hits are filtered by their distribution across taxonomic nodes until the most distant taxonomic node is determined as the ‘gene age’ (or removed as potential contamination), with the evolutionarily oldest genes assigned as phylostratum (PS) 1 and the youngest assigned as PS 8 in *F. serratus* and *F. distichus* and PS 11 in *Ectocarpus*. PS 7 corresponds to the origin of brown algae (complex multicellularity). For genes with more than one isoform, the age of the oldest isoform was used. Thus, after filtering lowly expressed genes across all samples (TPM<2) and potential contaminations, we obtained expression and evolutionary data for 8,291 genes in *F. serratus*, 7,907 genes in *F. distichus* and 11,571 genes in *Ectocarpus*. Using myTAI v1.0.1.9000 (Drost et al. 2018), TAI was calculated for each stage (*TAI_s_*) as follows,

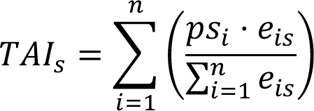

where *рs_i_* denotes the relative gene age (phylostratum) for a given gene *i*. The term *e_is_* denotes the expression level of a given gene *i* at developmental stage *s* and *n* denotes the total number of genes.

The expression level was captured using TPM values, since we are quantifying the relative abundance of mRNA molecules per gene rather than the count of sequencing fragments. To test the stability of the TAI profiles and reduce the variance in the highly expressed genes (Piasecka et al. 2013), we performed several RNA-seq data transformations on the expression matrices: square-root transformation (used for the main figures), log transformation with a pseudo-count of 1 (i.e., log2(TPM+1)), ‘regularised log’ transformation (i.e., rlog) (Love et al. 2014), and rank transformation (i.e., genes were ranked by level of expression at each stage). To reduce potential outliers, the median abundance value of replicates was chosen to represent the expression level (*e_is_*).

The statistical significance of the resulting profiles was assessed using non-parametric permutation tests (flat line test and reductive hourglass test), using the `FlatLineTest()` and `ReductiveHourglassTest()` functions implemented in myTAI (Drost et al. 2018). We defined ‘early’ stages as S1-2, ‘mid’ as S3-4, and ‘late’ as S5 in *F. serratus*, and ‘early’ stages as S1-4, ‘mid’ as S4.5, and ‘late’ as S5 in *F. distichus*, due to differences in developmental stage correspondence. These tests, including those for sex differences, were performed with 50,000 permutations.

For the pTAI analysis, we used the function `pMatrix()` implemented in myTAI (Drost et al. 2018), which calculates the contribution of each gene to the TAI at each stage by multiplying the phylostratum of each gene by its expression level divided by the sum of expression of all genes, i.e.

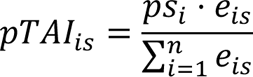

where, like TAI, *рs_i_* denotes the relative gene age (phylostratum) for a given gene *i* and *e_is_*denotes the expression level of a given gene *i* at developmental stage *s* and *n* denotes the total number of genes. The elbow method was used to identify 500 genes with the highest TAI contribution in each developmental stage; genes driving the TAI value across all unicellular or multicellular stages were inferred via intersection. For consistency with the main TAI analyses, square-root transformation was applied before the pTAI analysis.

### Transcriptome specificity index

To investigate whether the transcriptome at the waist of the hourglass is composed of broadly expressed genes compared to other stages, we first indexed each gene by its relative expression specificity/breadth across development using *tau* (Yanai et al. 2005; Kryuchkova-Mostacci and Robinson-Rechavi 2017), i.e.

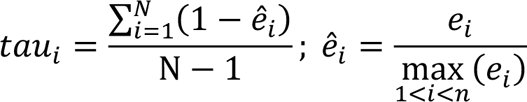

where *N* is the number of stages, *ê_i_* is the expression level of a given gene *i* normalised by the maximal expression value. A lower *tau* indicates low stage-specificity (in other words, broad expression), and *vice-versa*. The resulting *tau* values across all genes are stratified into deciles (tau-stratum), which enables analogous comparisons to TAI. It should be noted that PS and *tau* are not correlated (Kendall’s τ ≈ 0.05 in both *Fucus* species), indicating that these metrics capture independent signals. In contrast to the TAI, the TSI captures the average expression specificity/breadth of a given transcriptome, weighted by the expression level of each gene, i.e.

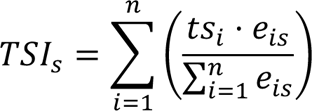

where *ts_i_* denotes the relative expression specificity/breadth (tau-stratum) for a given gene *i*, *e_is_*denotes the expression level of a given gene *i* at developmental stage *s* and *n* denotes the number of genes. The median abundance of replicates was chosen to represent the expression level (*e_is_*). Existing functions in myTAI were repurposed for this analysis, including non-parametric permutation tests (50,000 permutations) to assess the statistical significance of the resulting profiles.

### Transcriptome divergence index

To explore whether unicellular stages in *Ectocarpus* not only exhibited a young transcriptome, but also genes under relaxed purifying selection, we computed the TDI. In contrast to the TAI, the TDI captures the average gene selective pressure (divergence-stratum; based on the dNdS ratio) of a given transcriptome, weighted by the expression level of each gene. The divergence-stratum of each gene in *Ectocarpus* was inferred from dNdS ratios using orthologr (Drost et al. 2015). In brief, one-to-one orthologues were inferred between *Ectocarpus* sp. 7 and *Ectocarpus subulatus* (Dittami et al. 2020), using best reciprocal hits, and the dNdS ratio was computed using the default “Comeron” estimation method. Importantly, >99% of one-to-one orthologue comparisons fell below the dNdS ratio of 1, indicating that we are quantifying the degree of purifying selection. Next, the resulting dNdS ratios across all genes are stratified into deciles, with the scale ranging from 1 (strong purifying selection) to 10 (weakest purifying selection). *Fucus* species were precluded from this analysis due to the short divergence time between *F. serratus* and *F. distichus*, which meant that more than 10% of genes having dNdS of 0. For genes with more than one isoform, the divergence-stratum of the oldest isoform was used.

Using myTAI (Drost et al. 2018), we calculated the TDI for each stage (*TDI*) as follows,

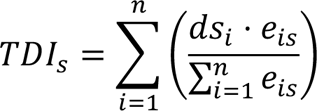

where *ds_i_* denotes the relative divergence level (divergence-stratum) for a given gene *i*. The term *e_is_* denotes the expression level of a given gene *i* at developmental stage *s*. The median abundance of replicates was chosen to represent the expression level (*e_is_*). The statistical significance of the resulting profiles was assessed using non-parametric permutation tests implemented in myTAI with 50,000 permutations.

### Distance/similarity-based transcriptome comparison

To quantify the overall distance/similarity between the transcriptomes of embryo stages in the two *Fucus* species, we computed the Pearson correlation, Spearman correlation, Manhattan distance and Jensen-Shannon distance (JSD) metric. Several metrics were employed owing to issues of calculating distance in high-dimensional data (Aggarwal et al. 2001). To compare expression levels between species, we compared the expression levels (i.e., abundance) of orthogroups (sets of orthologues and paralogues) using OrthoFinder v2.5.4 (Emms and Kelly 2019), treating genes as isoforms and orthogroups as genes when importing the RNA-seq data using tximport v1.26.1 (Soneson et al. 2016). Orthogroups were used rather than one-to-one orthologues (inferred through procedures such as best reciprocal hit), since orthogroups also capture the expression profile of in-paralogues, thereby covering more genes in the genome. The abundance data was transformed using rlog (Love et al. 2014). Correlation matrices were computed using `cor()` from the stats package in R (R Core Team 2022), while Manhattan and Jensen-Shannon Distance (JSD) metrics were computed using the R package philentropy v0.7.0 (Drost 2018).

### Enrichment analyses

GO enrichment analysis was performed on genes contributing most to *Ectocarpus* TAI using Fisher’s exact test statistics with the parent-child algorithm as implemented in TopGO (Grossmann et al. 2007; Alexa and Rahnenfuhrer 2023). Statistical tests and significance levels are indicated in the text and figure legends.

## Author contributions

JSL: Investigation (equal); Formal analysis (lead); Visualization (lead), Writing – original draft (lead)

MZ: Investigation (equal); Methodology (supporting)

RL: Investigation (equal), Methodology (supporting); Visualization (supporting)

H-GD: Conceptualization (equal); Funding acquisition (supporting); Data curation (equal); Methodology (lead); Visualization (supporting); Project administration (equal); Supervision (equal); Writing – original draft (equal); Writing – review and editing (supporting).

SMC: Conceptualization (lead); Funding acquisition (lead); Methodology (equal); Project administration (lead); Supervision (equal); Visualization (supporting); Writing – original draft (supporting); Writing – review and editing (lead).

## Acknowledgements

We thank Claudia Martinho, Daniel Liesner, Agnieszka Lipinska, James Tan Shen Yi and Josue Barrera-Redondo for insightful discussions. We also thank Nikola Kalábová for implementing multithreaded permutation tests in myTAI and Detlef Weigel for support and sponsorship of H-GD. This work was supported by the MPG, the ERC (grant n. 864038 to SMC) and the BMBF-funded de.NBI Cloud within the German Network for Bioinformatics Infrastructure (de.NBI) (031A532B, 031A533A, 031A533B, 031A534A, 031A535A, 031A537A, 031A537B, 031A537C, 031A537D, 031A538A). SMC is supported by the Moore Foundation (GBMF11489) and the Bettencourt-Schuller Foundation. Computations were also performed in the MPCDF Cobra supercomputer in Garching, Germany and the cluster of the Max Planck Campus in Tübingen, Germany. JSL thanks the International Max Planck Research School ‘From Molecules to Organisms’.

Supplementary Information is available for this paper.

Reprints and permissions information is available at www.nature.com/reprints.

## Data availability

Data accession codes are available in **Table S1**. Correspondence and requests for materials should be addressed to (research and materials) and (research and methods)

## Code availability

The analysis code is available at https://github.com/LotharukpongJS/hourglass_brownalga.

## Supplementary figures

**Figure S1.**
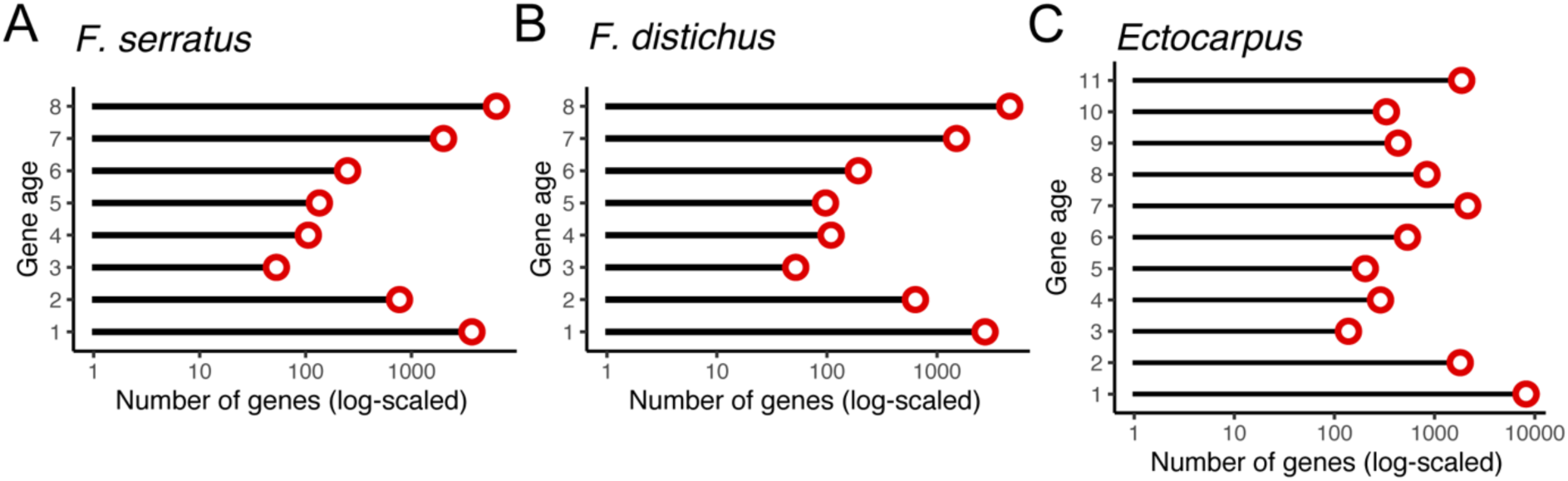
Summary statistics for the phylostratigraphic maps of *F. serratus*, *F. distichus*, and *Ectocarpus* (species 7). Taxonomic ranks and the number of genes assigned to each rank (phylostratum - gene age ranks) in brown algae: (A) *F. serratus*, (B) *F. distichus* and (C) *Ectocarpus*.

**Figure S2.**
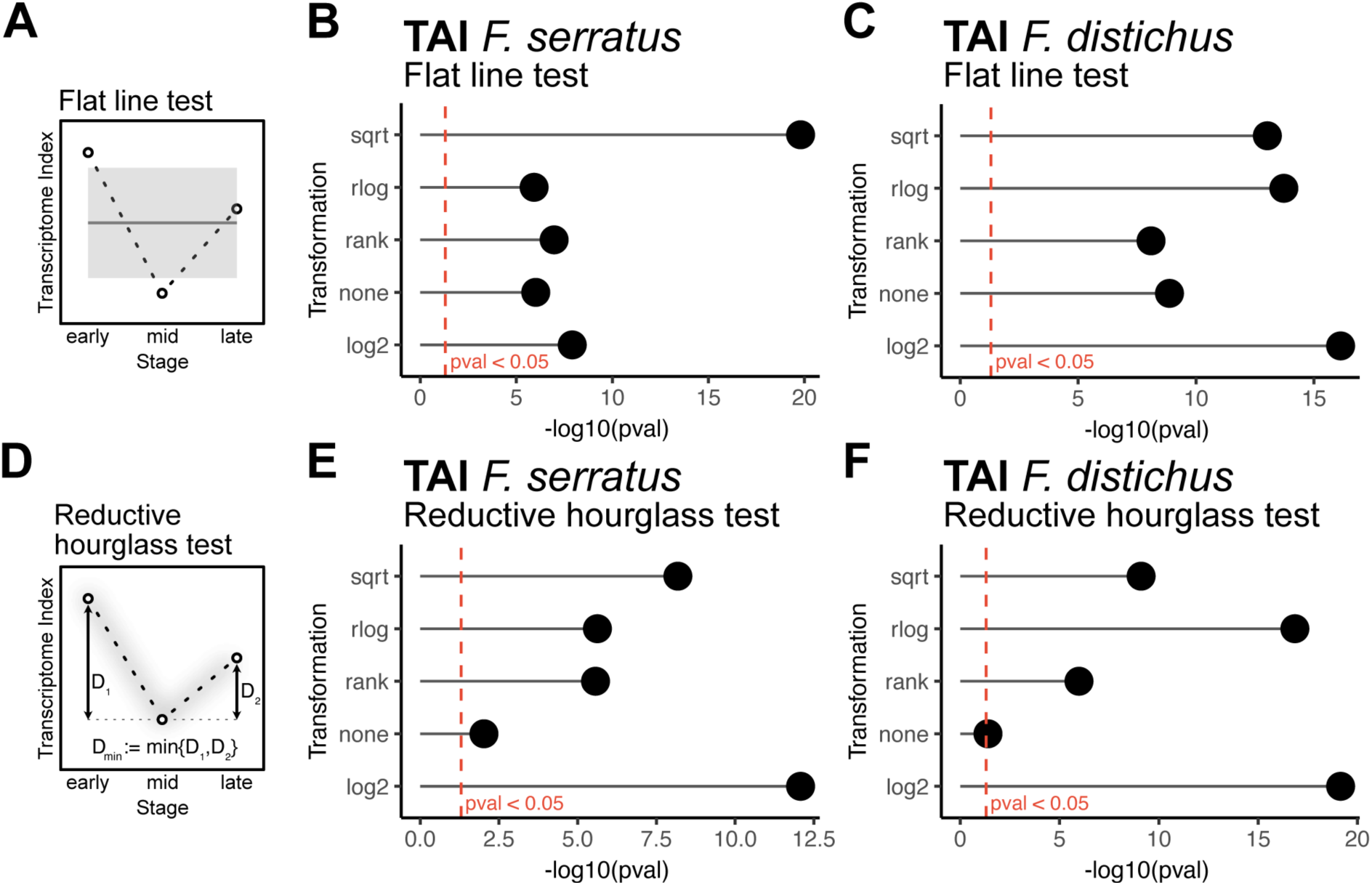
Permutation tests on the evolutionary transcriptome patterns in *Fucus* development under various RNA-seq transformations. (A) The flat line test evaluates any significant deviation of an observed evolutionary transcriptome pattern from a flat line. Flat line tests in (B) *F. serratus* and (C) *F. distichus*, under various RNA-seq transformations. (D) The reductive hourglass test evaluates whether an observed evolutionary transcriptome pattern follows an hourglass (high-low-high) pattern, based on the statistical significance of the observed minimum difference (D_min_) between the ‘mid’ module and the ‘early’ (D_1_) or ‘late’ (D_2_) modules. Reductive hourglass tests in (E) *F. serratus* and (F) *F. distichus*, under various RNA-seq transformations. All tests were conducted with 50,000 permutations.

**Figure S3.**
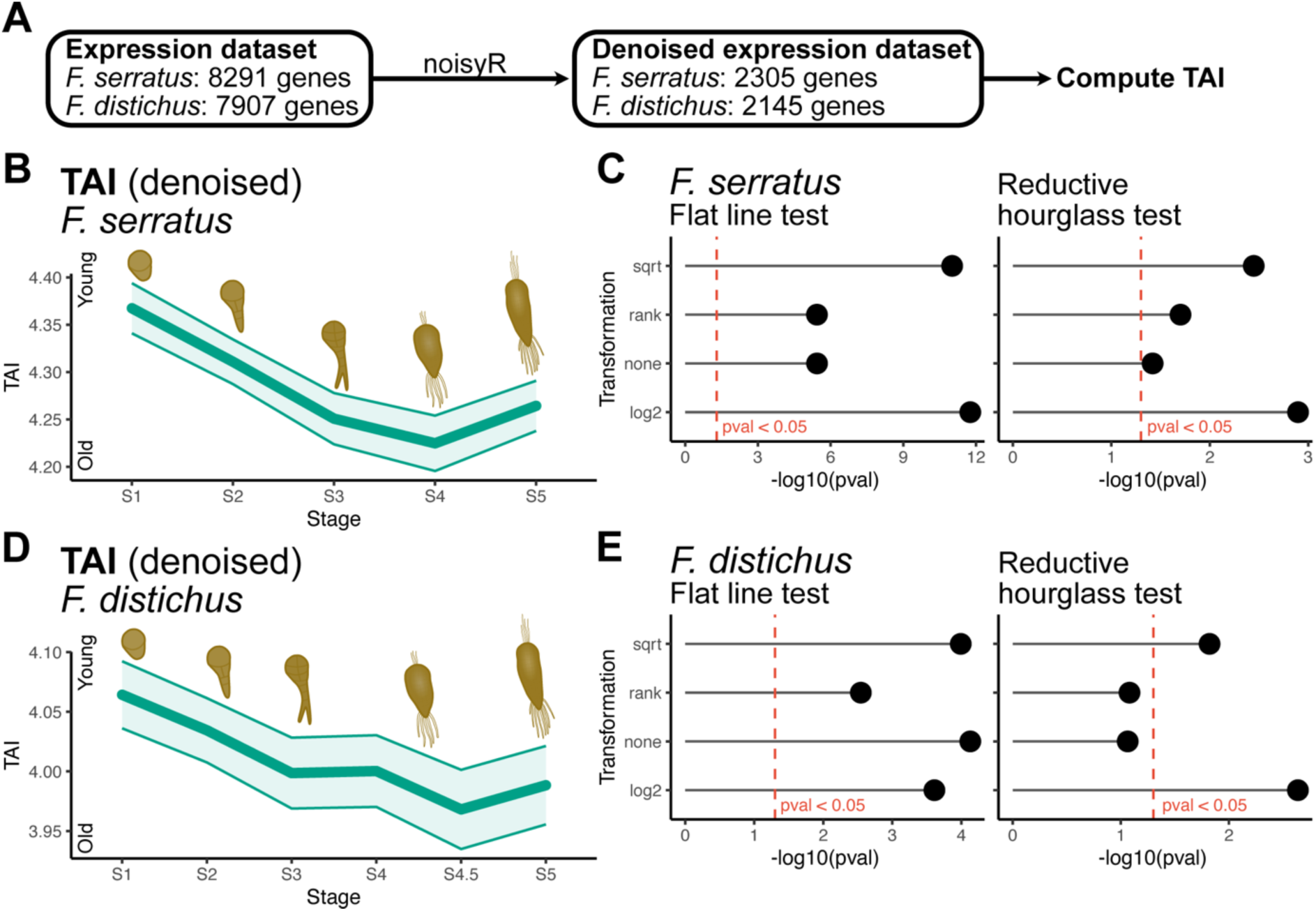
Transcriptome age patterns in *Fucus* development using a denoised dataset. (A) Overview of removal method for genes with ‘noisy’ expression profile using noisyR. (B) TAI across embryogenesis in *F. serratus* using a denoised dataset. (C) Flat line and reductive hourglass permutation tests in *F. serratus* under various RNA-seq transformations. (D) TAI across embryogenesis in *F. distichus* using a denoised dataset. (E) Flat line and reductive hourglass permutation tests in *F. distichus* under various RNA-seq transformations. 50,000 bootstraps were used to compute the standard deviation in (B) and (D). Permutation tests were performed with 50,000 permutations.

**Figure S4.**
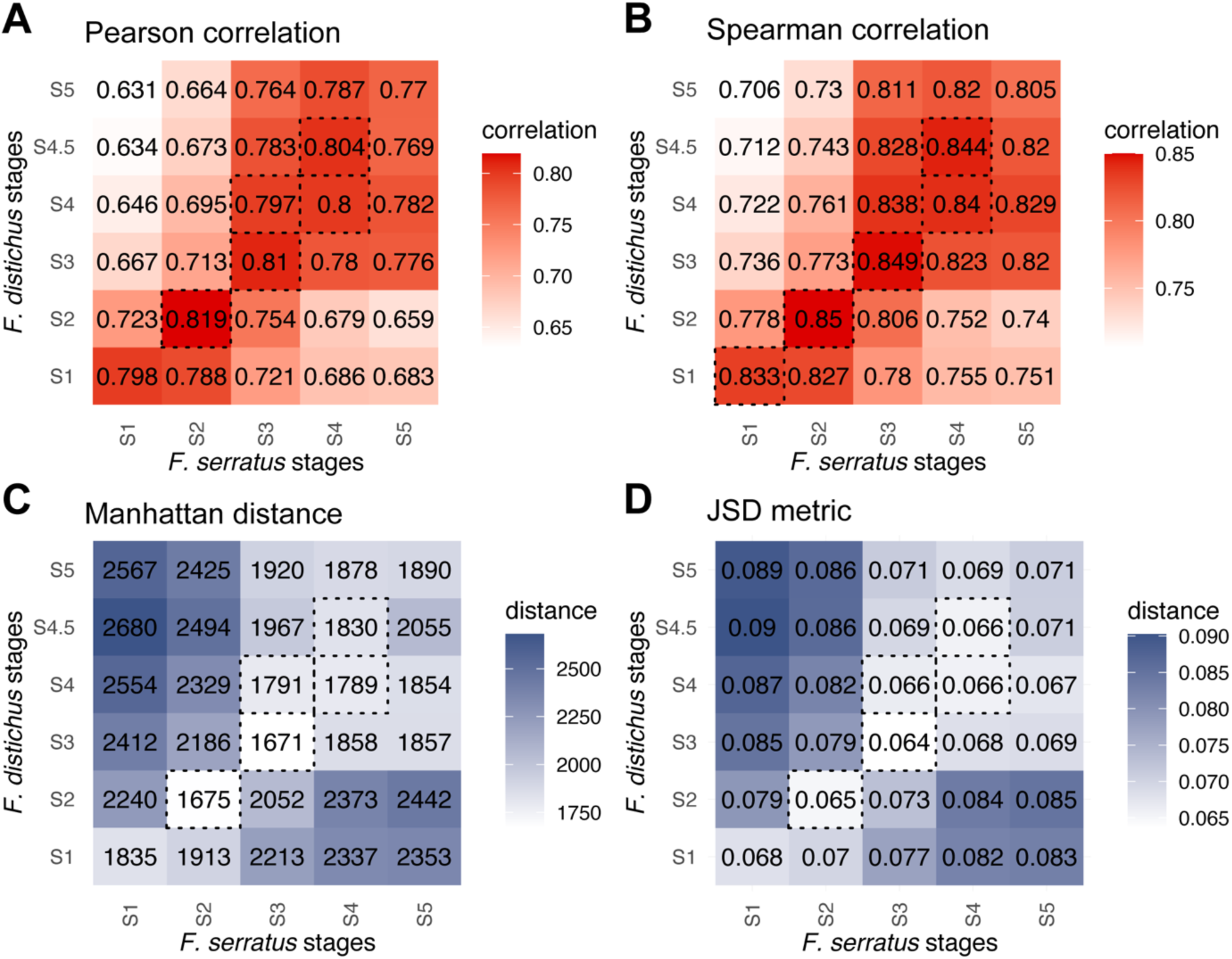
Correspondence between embryo stages of *Fucus* species using expression distance/similarity. Expression levels of shared orthogroups were compared rather than one-to-one orthologues to mitigate the effect of in-paralogues. Shown are the median (A) Pearson correlation, (B) Spearman correlation, (C) Manhattan distance and (D) Jensen-Shannon Distance (JSD) metric between each embryonic stage under the rlog transformation. Dashed boxes indicate the five most similar/closest comparisons between the two closely related species.

**Figure S5.**
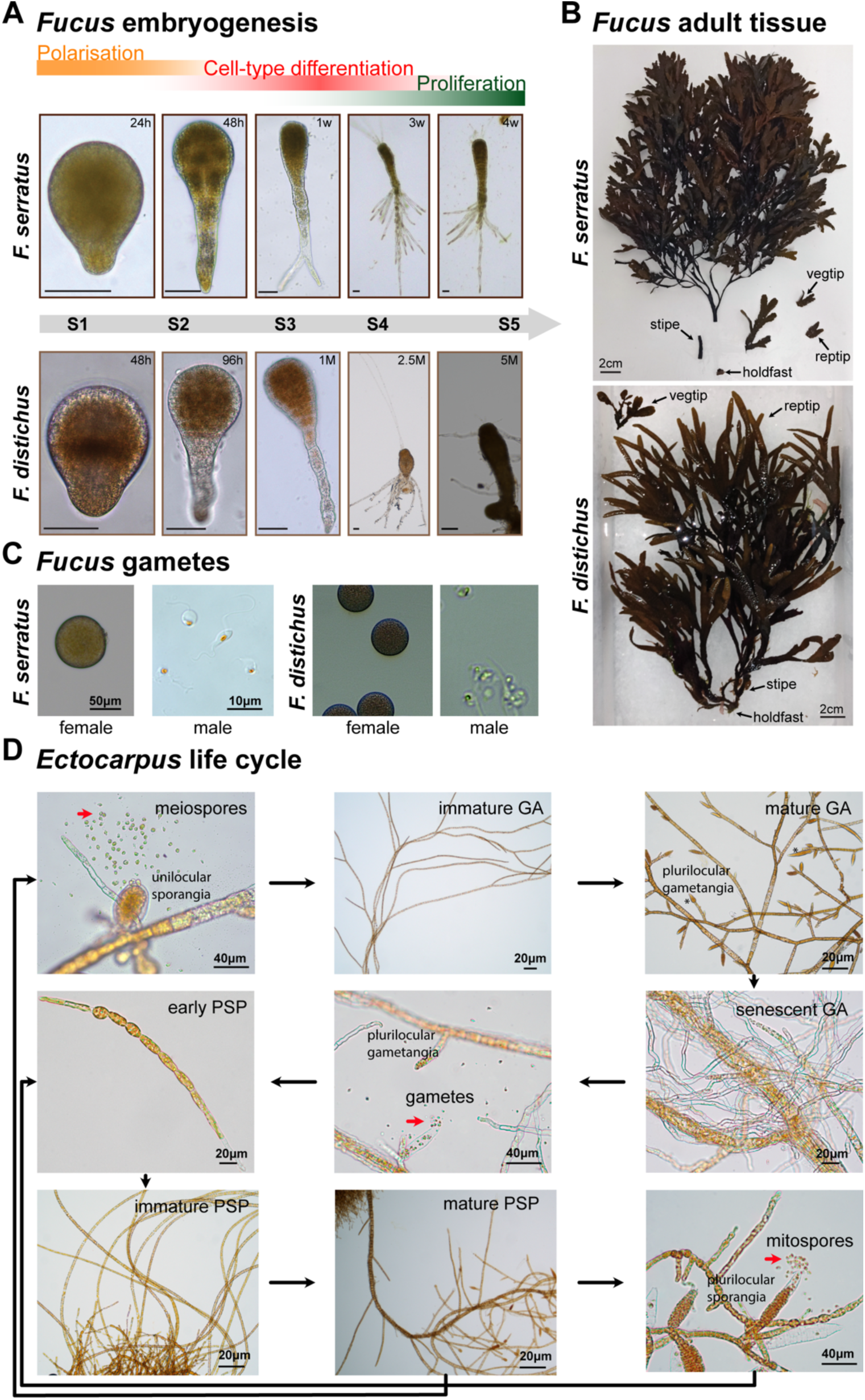
Images of life cycle stages of *Fucus* and *Ectocarpus*. (A) Embryo stages of *F. serratus* and *F. distichus*. In both species, the embryos go through phases of polarisation, cell-type differentiation and proliferation of established cell types. (B) Adult tissues of *F. serratus* and *F. distichus*. (C) Gametes of *F. serratus* and *F. distichus*. (D) *Ectocarpus* life cycle stages. Black arrows show development. Red arrows point to unicells.

**Figure S6.**
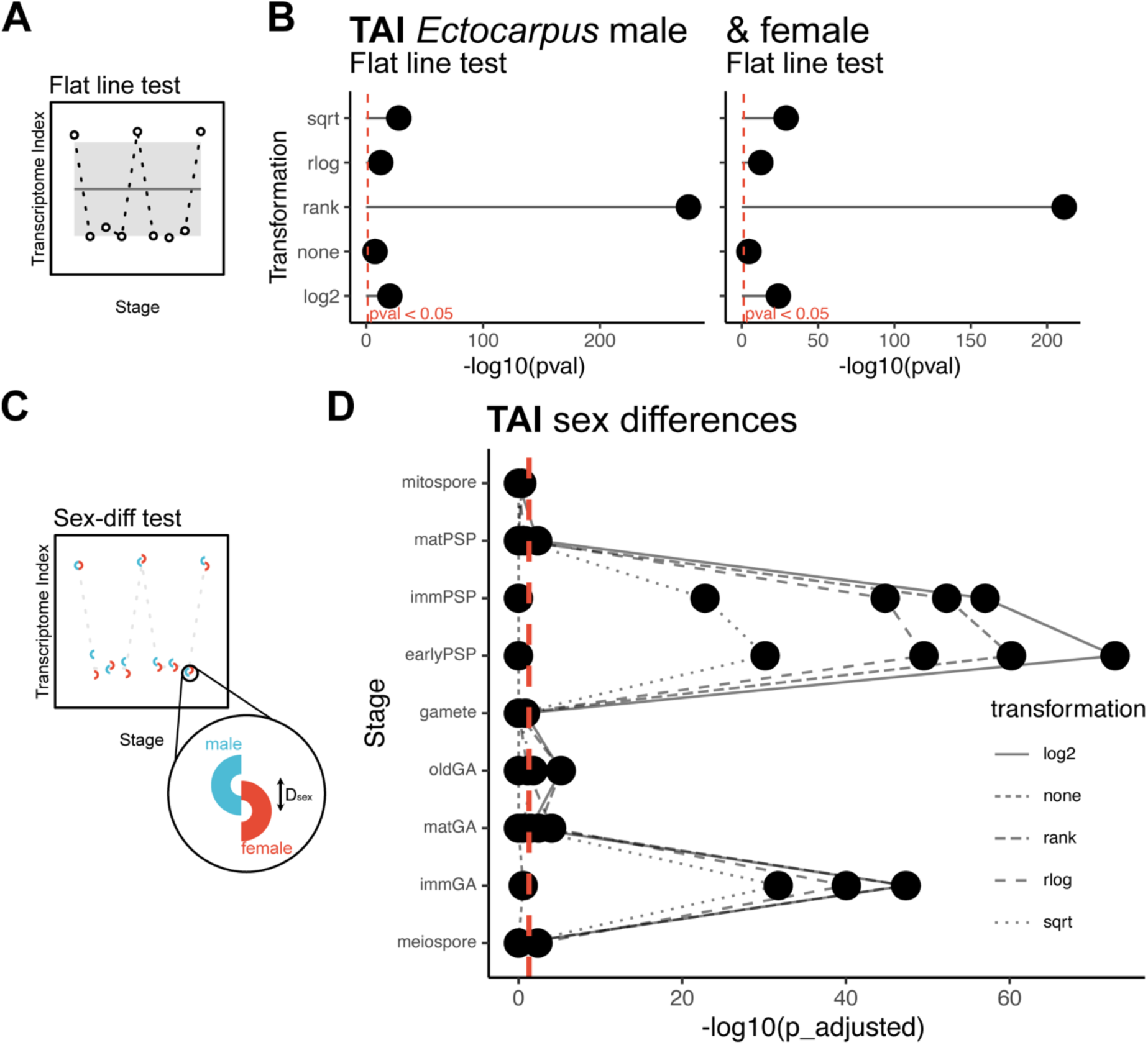
Permutation tests on the evolutionary transcriptome patterns in *Ectocarpus*. (A) The flat line test evaluates any significant deviation of an observed evolutionary transcriptome pattern from a flat line. (B) Flat line tests in male and female *Ectocarpus*, under various RNA-seq transformations. All profiles were statistically significant. (C) The sex difference permutation test evaluates sex differences (D_sex_) in the evolutionary transcriptome at a given stage. (D) Differences in TAI between sexes at each developmental stage in *Ectocarpus*, under various RNA-seq transformations. The resulting p-values were adjusted using Bonferroni correction. All tests were conducted with 50,000 permutations.

**Figure S7.**
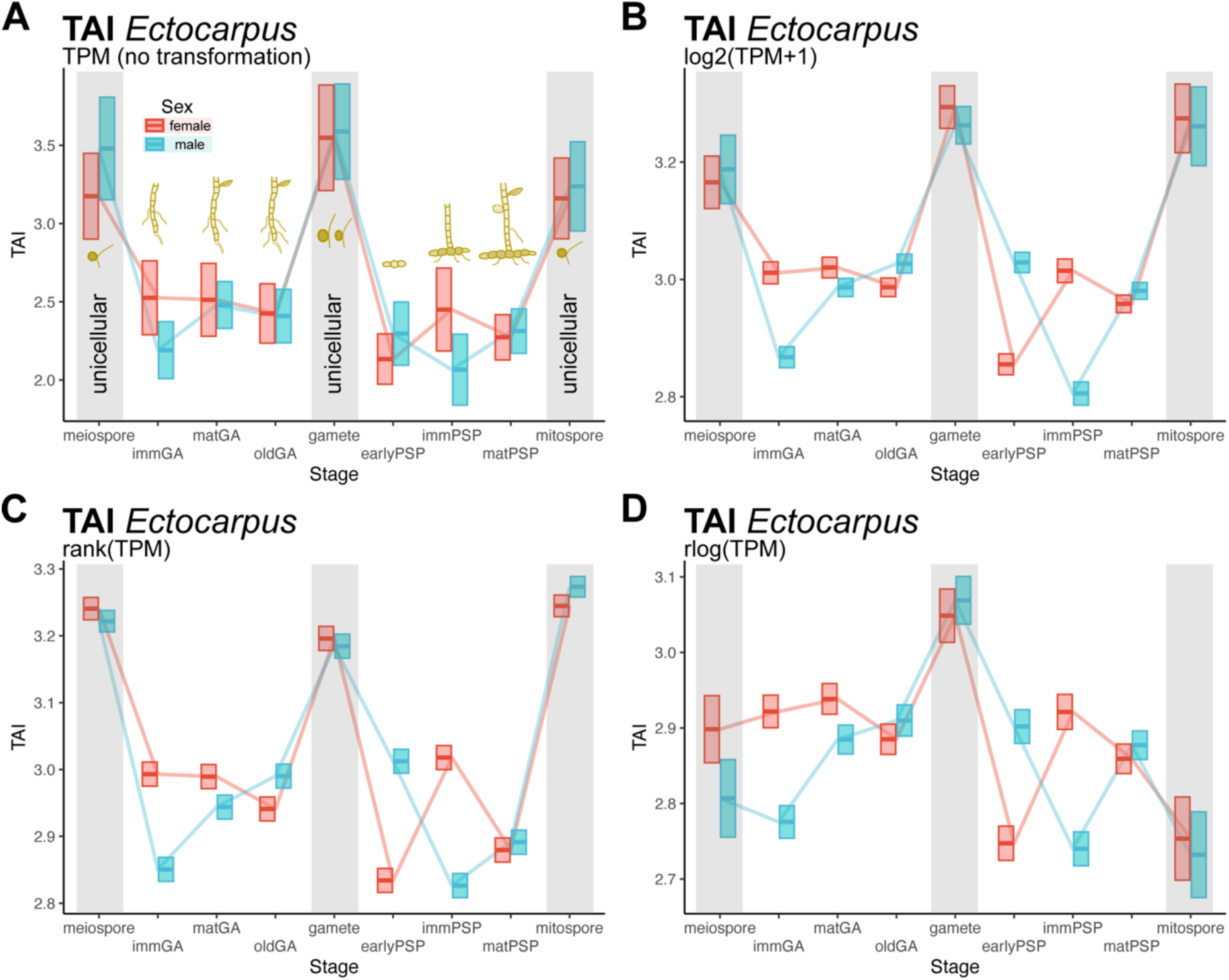
TAI profile across life cycle stages of *Ectocarpus*, under different RNA-seq transformations. TAI was computed after applying different RNA-seq transformations: (A) raw TPM (none), (B) log2(TPM+1), (C) rank, and (D) regularised log. A lower TAI marks a transcriptome composed of evolutionarily older genes and *vice versa*. Blue lines represent male stages while red lines represent female stages. The upper and lower limits of the boxes demarcate the standard deviation, based on 50,000 bootstraps.

**Figure S8.**
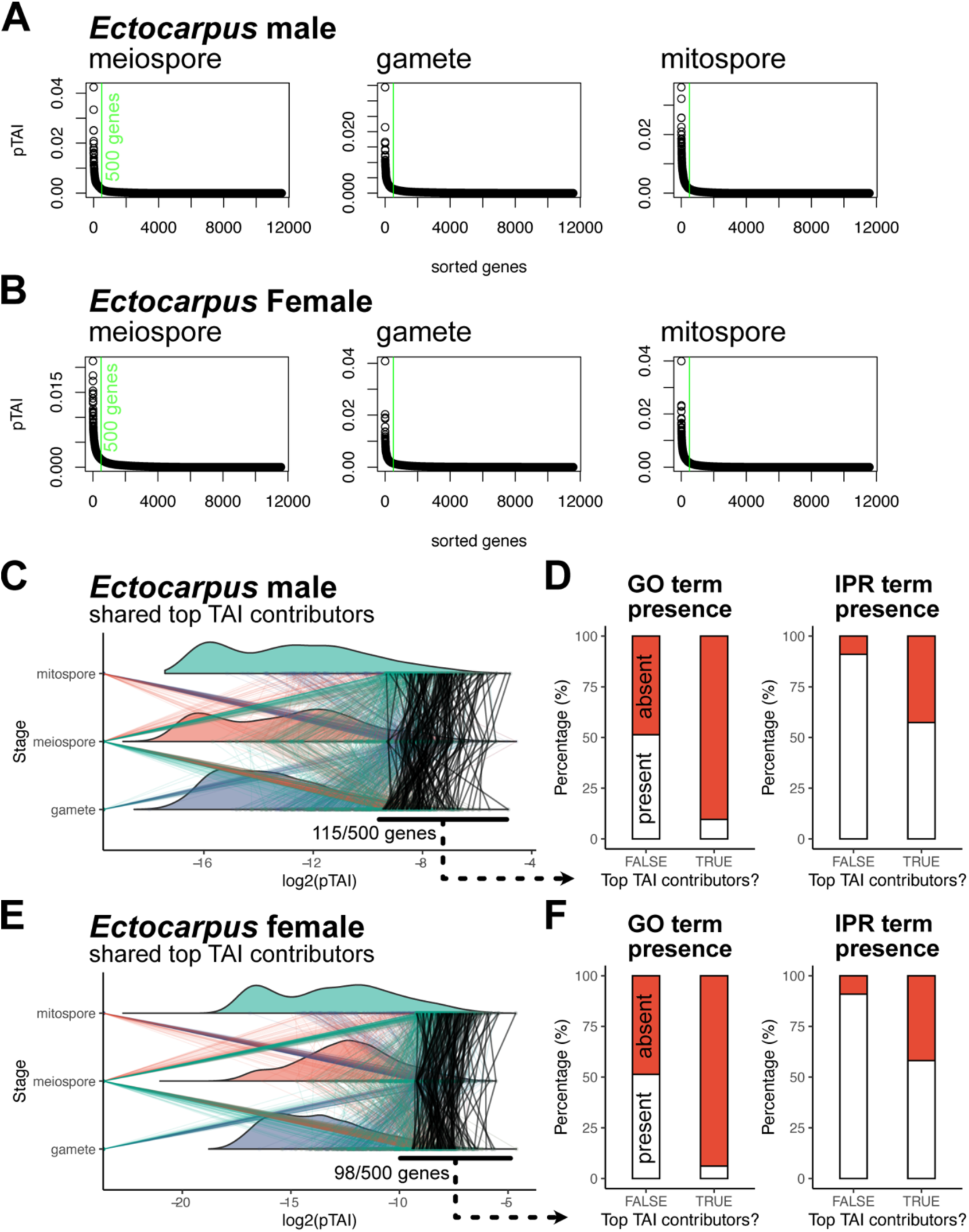
pTAI in unicellular stages of *Ectocarpus*. Genes ranked by their pTAI across each unicellular stage of (A) male and (B) female *Ectocarpus*. The green line indicates the top 500 gene cutoff based on the elbow method, i.e., genes that explain most of the overall TAI (‘top contributors’). (C) Shared top contributors in unicellular stages of male *Ectocarpus*. The coloured lines denote the pTAI value of a given top-contributor gene in one unicellular stage mapped across other unicellular stages. Shared top contributors across all stages are denoted by black lines. (D) The presence/absence of gene ontology (GO) and interproscan (IPR) domain terms in the shared top contributors in unicellular stages of *Ectocarpus*. The same analysis done on female *Ectocarpus* are shown in (E & F). For visual clarity, the pTAI values were log-transformed.

**Figure S9.**
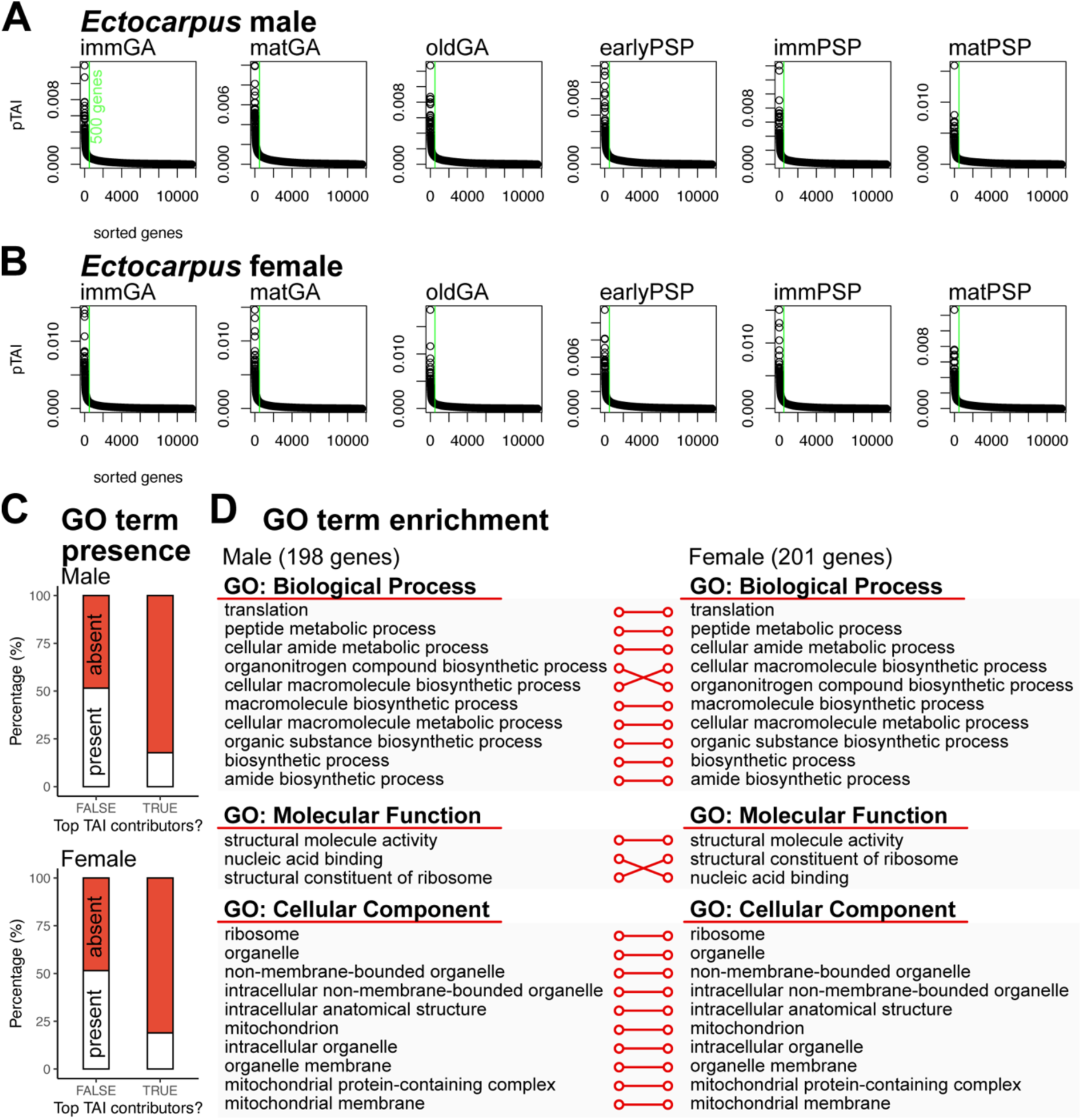
pTAI in multicellular stages of *Ectocarpus*. Genes ranked by their pTAI across each multicellular stage of (A) male and (B) female *Ectocarpus*. The green line indicates the top 500 gene cutoff based on the elbow method, i.e., genes that explain most of the overall TAI (‘top contributors’). (C) The presence/absence of gene ontology (GO) terms in the shared top contributors in multicellular stages of *Ectocarpus* (198 in males and 201 in females). (D) The relationship between ten most significant GO terms ranked by p-values for each GO category between male and female *Ectocarpus*. Significance is defined by padj < 0.1.

## Supplementary tables

**Table S1.** Sample list and sequencing data details including accessions.

**Table S2.** Gene age and expression data per developmental stage (‘phyloexpression set’ format), separated into six sheets. (1) *F. serratus*, (2) *F. distichus*, (3) *Ectocarpus* male strain EC32m, (4) *Ectocarpus* female strain EC25f, (5) *F. serratus* denoised, (6) *F. distichus* denoised.

**Table S3.** *tau* values per gene, separated into two sheets. (1) *F. serratus*, (2) *F. distichus*.

**Table S4.** dNdS values per gene in *Ectocarpus* (sp. 7), detailing the pairwise alignment match (reciprocal best hit) with *E. subulatus*, alignment scores, dN values, dS values and dNdS values.

**Table S5.** GO terms from pTAI analysis, separated into four sheets. (1) *Ectocarpus* unicellular male (note: all non-significant), (2) *Ectocarpus* unicellular female (note: all non-significant), (3) *Ectocarpus* multicellular male, (4) *Ectocarpus* multicellular female.

